# Human mitochondrial protein complexes revealed by large-scale coevolution analysis and deep learning-based structure modeling

**DOI:** 10.1101/2021.09.14.460228

**Authors:** Jimin Pei, Jing Zhang, Qian Cong

**Affiliations:** McDermott Center for Human Growth and Development, University of Texas Southwestern, Medical Center at Dallas, 6001 Forest Park Rd., Dallas, TX, USA, Texas, U.S.A. 75390

**Keywords:** mitochondrial proteins, coevolution analysis, protein-protein interactions, RoseTTAFold, AlphaFold2

## Abstract

Recent development of deep-learning methods has led to a breakthrough in the prediction accuracy of 3-dimensional protein structures. Extending these methods to protein pairs is expected to allow large-scale detection of protein-protein interactions and modeling protein complexes at the proteome level. We applied RoseTTAFold and AlphaFold2, two of the latest deep-learning methods for structure predictions, to analyze coevolution of human proteins residing in mitochondria, an organelle of vital importance in many cellular processes including energy production, metabolism, cell death, and antiviral response. Variations in mitochondrial proteins have been linked to a plethora of human diseases and genetic conditions. RoseTTAFold, with high computational speed, was used to predict the coevolution of about 95% of mitochondrial protein pairs. Top-ranked pairs were further subject to the modeling of the complex structures by AlphaFold2, which also produced contact probability with high precision and in many cases consistent with RoseTTAFold. Most of the top ranked pairs with high contact probability were supported by known protein-protein interactions and/or similarities to experimental structural complexes. For high-scoring pairs without experimental complex structures, our coevolution analyses and structural models shed light on the details of their interfaces, including CHCHD4-AIFM1, MTERF3-TRUB2, FMC1-ATPAF2, ECSIT-NDUFAF1 and COQ7-COQ9, among others. We also identified novel PPIs (PYURF-NDUFAF5, LYRM1-MTRF1L and COA8-COX10) for several proteins without experimentally characterized interaction partners, leading to predictions of their molecular functions and the biological processes they are involved in.

## Introduction

Recent advances in Deep-Learning (DL) techniques for structure prediction based on protein-sequence alignments have led to a breakthrough in structural genomics. These state-of-the-art methods, which are now sensitive enough to work with shallow alignments, can predict protein 3D structure to atomic accuracy [1-3]. The methods can be applied on the whole-proteome scale [4], with the 3D structures for almost all human proteins being recently “determined” by DeepMind using Alphafold2. While the quality of these 3D structures remains to be validated by the scientific community, they unquestionably facilitate functional characterization of these proteins and interpretation of disease-causing mutations in them [4].

One of the next research directions where these DL methods can transform is to determine protein-protein interactomes. Using statistical analyses of coevolutionary signal between residues from different proteins, we have shown that the interface between interacting proteins in Bacteria can be accurately predicted from alignments of diverse sequences on the proteome scale [5]. This type of *in silico* protein-protein interaction screen appears to be more accurate than large-scale experimental screens such as yeast-two-hybrid and affinity-purification-mass-spectrometry [5]. The application of similar methods to human sequences were hindered by limitations in sensitivity for shallower Eukaryotic sequence alignments. However, we expect the increased performance of current DL methods will enable proteome-wide screen of protein-protein interaction (PPI) in human.

Mitochondria are dynamic membrane-bound organelles found in almost all eukaryotic cells. They carry a variety of essential cellular functions ranging from energy production and metabolism to regulation of cell death and immune response [6-10]. Mitochondria evolved from an endosymbiotic relationship between ancestral bacteria and eukaryotes [11]. While these organelles maintain their own genetic materials and transcriptional and translational machineries, most proteins targeted to mitochondria are now encoded in the nuclear genome. Technology advances in genomics, transcriptomics, proteomics, and metabolomics have extended the repertoire of mitochondrion-associated proteins, identified their interaction partners, and expanded the knowledge about their functions [12-16]. The MitoCarta3.0 database has summarized 1136 mitochondrial proteins in human and classified them according to their pathways [17]. These mitochondrial proteins are located in four compartments: the mitochondrial outer membrane (MOM), the intermembrane space (IMS), the mitochondrial inner membrane (MIM), and the matrix. Characterizations of mitochondrial proteins and their complexes are crucial to our understanding of how mitochondria regulate various cellular activities and the mechanisms of a diverse range of mitochondrion-related diseases [18].

As a pilot study, we applied RoseTTAFold [1] and Alphafold2 [2] to identify interacting protein pairs and predict their interfaces among human mitochondrial proteins. We showed that a combination of these methods can identify interacting partners and model protein complexes on a large-scale with high precision. With these methods, we captured many known PPIs that were supported by multiple experiments and/or protein complex structures. In addition, we obtained detailed information about the interfaces of known PPIs without experimentally determined complex structures and identified novel PPIs for several proteins without experimentally characterized interaction partners. These findings provided structural insight into the mechanisms of some disease-causing mutations and functional predictions of several poorly characterized proteins.

## Results and discussion

### In silico screen can identify interacting proteins and predict their interface at high precision

We designed a computational procedure (flowchart in Figure 1A, see Materials and methods) to perform *in silico* PPI screen for 605099 pairs of 1118 mitochondrial proteins from the MitoCarta3.0 database [17] using RoseTTAFold [1] and AphaFold2 [2]. About 5% of mitochondrial protein pairs were excluded from this study because their combined lengths were beyond our GPU’s compute capability. We used the best inter-residue score (see Materials and methods) between a pair of proteins to indicate their interaction confidence. Precision-recall curves based on true PPIs (positive controls) and random protein pairs (negative controls, see Materials and methods) evaluate the performance of these methods. The RoseTTAFold 2-track model, a fast version of RoseTTAFold without 3D representation of the proteins inside the model, is remarkably better than Direct Coupling Analysis (DCA), a statistical method we used previously in coevolution-based Bacteria PPI screens [5] (Figure 1B). The area under the precision-recall curve scores (AUC) for DCA (0.014) is much lower than the AUC for RoseTTAFold (0.113). In addition, we observed that residues in the mitochondria-targeting peptides of different proteins can show high contact probability, although they are not expected to be involved in protein-protein interface as they are cleaved after being transferred to mitochondria. Excluding the mitochondrion-targeting peptides before computing the highest inter-residue contact probability for a pair of proteins further improved the accuracy of the screen (Figure. 1C) (AUC improved from 0.113 to 0.132).

**Figure 1.**
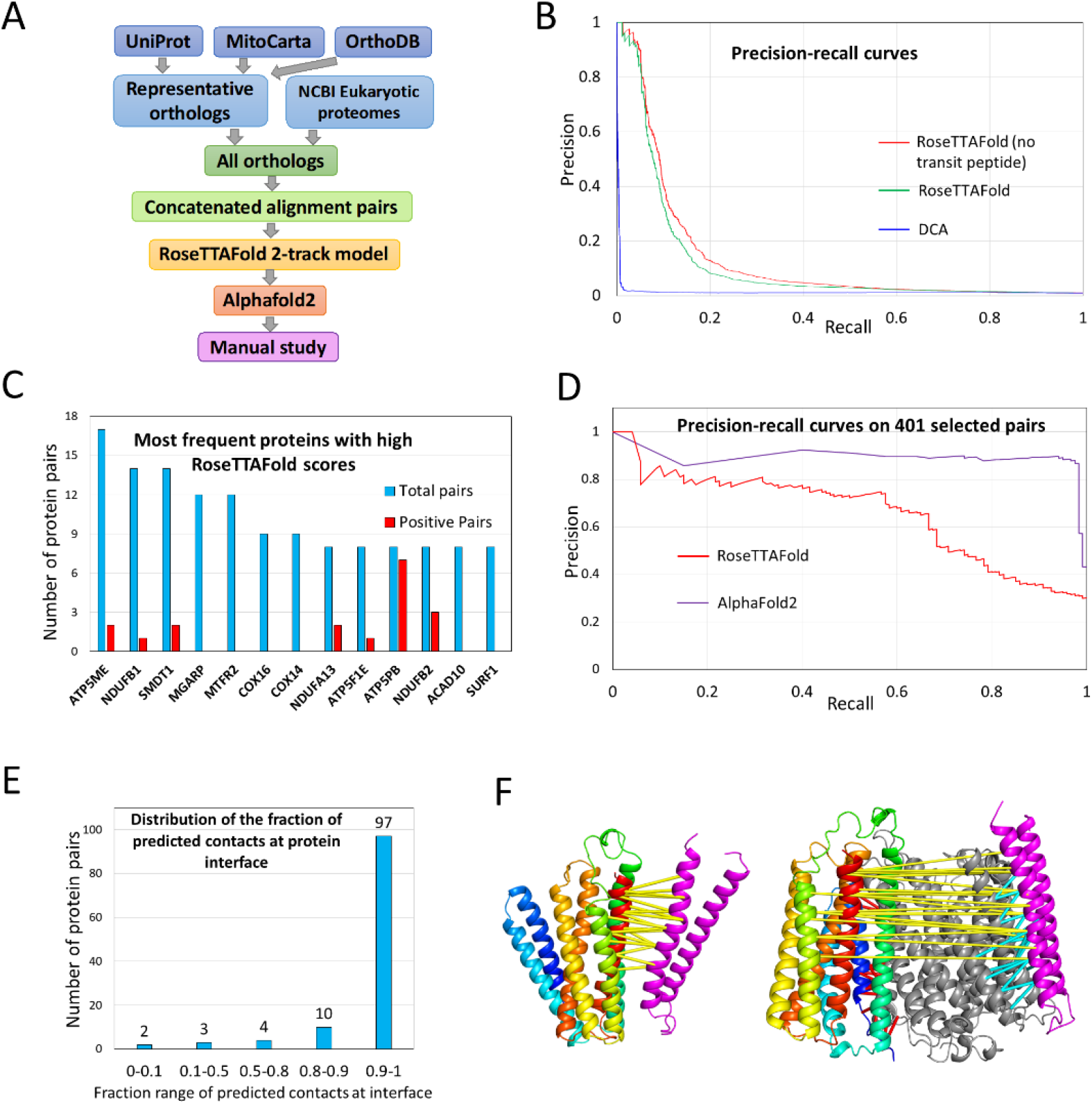
**A**. A flowchart of the procedure used in this study to perform PPI prediction and structural modeling. **B**. The precision-recall curves of three methods in detecting positive PPIs of mitochondrial protein pairs. DCA - Direct Coupling Analysis method, RoseTTAFold (no transit peptide) is the RoseTTAFold method used on residue pairs excluding those involving any residue in mitochondrial transit peptides. **C**. Counts of high-scoring interacting partners (contact probability score >0.9) (blue bars) for proteins with the highest number of such partners. The red bars (if any) show the number of true positives among the top-scoring pairs. **D**. Precision-recall curves for RoseTTFold and AlphaFold2 on the 401 pairs of selected PPI predictions. **E**. Distribution of the fraction of predicted contacts that are mapped to experimentally determined complex interfaces. **F**. The left panel shows the AlphaFold2 model of MT-CO2 (magenta) and MT-CO3 (rainbow) complex, with the top 20 RoseTTAFold-predicted contact pairs shown as yellow lines connecting their C-α atoms. The right panel shows the AlphaFold2 model of the MT-CO1 (gray), MT-CO2 (magenta) and MT-CO3 (rainbow) complex. Top contact pairs are shown in yellow, cyan, and red lines for MT-CO2/MT-CO3, MT-CO1/MT-CO2 and MT-CO1/MT-CO3 pairs, respectively.

The majority of the top 100 pairs (70 out of 100) with the highest RoseTTAFold contact probability are PPIs supported by either experimental structural complexes or multiple incidences (≥ 3) in the BioGRID [19] database. They include 37 pairs involving mitochondrial ribosomal subunits [20, 21] and more than 10 pairs involving oxidative phosphorylation (OXPHOS) complex I (CI, NADH:ubiquinone oxidoreductase complex) subunits or assembly factors [22, 23]. Multiple PPIs were also found between the subunits and assembly factors of OXPHOS complex II (CII, succinate-Q oxidoreductase complex) [24], complex III (CIII, cytochrome bc1 complex), complex IV (CIV, ubiquinol-cytochrome c oxidase complex), and complex V (CV, ATP synthase complex) [23], respectively. Other protein pairs from the same complexes identified as top hits of our *in silico* screen include SUCLG1-SUCLG2 and SUCLG1-SUCLA2 from the succinyl-CoA synthetase complexes [25], PDHA1-PDHB and PDHA2-PDHB from the pyruvate dehydrogenase complex [26], ETFA-ETFB from the electron transfer flavoprotein complex [27], BCKDHA-BCKDHB from the branched-chain alpha-keto acid dehydrogenase complex [28], GATB-GATC and GATC-QRSL1 from the glutaminyl-tRNA amidotransferase complex [29], and TIMM9-TIMM10 from the translocase of the inner membrane (TIM) complex [30].

One known artifact of statistical methods for inter-protein coevolution analysis occurs for some proteins showing a high coevolution signal with many other proteins [5]. We observed the same artifact implementing RoseTTAFold on mitochondrial PPIs (Figure. 1C). 13 proteins exhibit a contact probability greater than 0.9 for 8 or more different interaction partners. Except for ATP5PB, interactions for most of these pairs are not supported by experiments (either experimental complex structures or BioGRID count ≥3) and could be false positives. To penalize such hubs of false positives, we adjusted the contact probability between a pair of proteins by a weight measuring each protein’s top contact probability with other proteins (see Materials and methods). We ranked the candidate PPIs by both the original contact probability and the adjusted contact score and selected 401 pairs as potential interacting partners (see Materials and methods, supplementary Table S1).

We modeled the 3D structure for these 401 pairs using AlphaFold2 [2]. Surprisingly, although AlphaFold2 was not trained to distinguish true PPIs from false ones, it exhibits remarkable performance on this task. It ranks the majority of the true positives among the 401 candidate PPIs higher than other pairs, achieving high precision (Figure 1D): 136 pairs among the 401 candidate PPIs had a top contact probability above 0.5 by Alphafold2, and 118 of them are considered true positives according to our criteria (supported by experimental structure or BioGRID count ≥ 3), indicating a precision of at least 87%. Those top hits not meeting our criteria could still be true PPIs, as some of them are supported by low-throughput experimental studies. Interesting examples are discussed in the following sections. At this level of precision, we recovered 118 of the 1563 true PPIs in the positive control set, indicating a recall of 7.5%.

It is important to note that applying AlphaFold2 to all pairs of mitochondrial proteins was computationally intractable (need ∼100,000 GPU hours), and reported performance was achieved by the sequential application of the fast RoseTTAFold 2-track model and AlphaFold2. We expect a better recall could be achieved by applying AlphaFold2 to all pairs of mitochondrial proteins. To probe the recall of AlphaFold2 for true PPIs, we applied it to all 473 pairs of mitochondrial proteins with confident experimental evidence (BioGRID reference count >= 3) for their interaction, and 120 pairs (25.4%) had top AlphaFold2 contact probability above 0.5 (Supplementary Table S2). We deposited the Alphafold2 models of these protein pairs at https://osf.io/g37mz/ The high accuracy of these AlphaFold2 models makes them a valuable resource to understand complex function, guide experimental design and explain disease mechanisms.

Out of the 401 interacting pairs we predicted *de novo* in our screen, 116 have known complex structure or close templates in the Protein Data Bank. We used these cases to evaluate the accuracy of residues contributing to the protein interface as suggested by pairwise RoseTTAFold scores. Most of these RoseTTAFold contacts show near perfect agreement with the experimental structures (Figure 1E). Only two pairs of proteins have low agreement between the contact residues predicted by RoseTTAFold and the interfaces defined by experimental complexes. One of the poorly predicted pairs is between MT-CO2 (COX2) and MT-CO3 (COX3), two mitochondrion-encoded CIV subunits. RoseTTAFold assigns high contact probability between the second transmembrane segment of MT-CO2 and the fourth and seventh transmembrane segments of COX3, in agreement with the AlphaFold2 structural model of MT-CO2 and MT-CO3 (Figure 1F, left panel). However, in experimental CIV structures, the transmembrane segments of MT-CO2 and MT-CO3 are separated by MT-CO1 and do not make contacts. We hypothesized that this wrong prediction was because we modeled a complex of MT-CO2 and MT-CO3 in the absence of MT-CO1. Therefore, we tried to model MT-CO1, MT-CO2, and MT-CO3 together by AlphaFold2, which generated a complex model in agreement with experimental structures (Figure 1F, right panel). The RoseTTAFold-predicted contacts between MT-CO1 and MT-CO2 (shown in red) and between MT-CO1 and MT-CO3 (shown in cyan) agree with this trimeric complex structure.

### Coevolution analysis revealed protein interaction interfaces for known protein complexes without experimental 3D structures

Several top-ranking protein pairs from our screen were found to have experimental support for their interactions but lack experimentally determined structures of the complexes. Our coevolution analysis supports the interactions between these protein pairs and provides detailed structural information on their interaction interfaces. Some of these pairs are described below.

#### CHCHD4 and AIFM1

AIFM1 (Apoptosis-Inducing Factor 1, Mitochondria, also named AIF) is a mitochondrial flavoprotein with multiple functions in both cell survival and cell death. It is essential for maintaining mitochondrial integrity in healthy cells and plays an important role in the apoptosis pathway [31, 32]. Anchored to the MIM by an N-terminal transmembrane segment, the majority of AIFM1localizes to the IMS. AIFM1 exhibits FAD-dependent NADH oxidase activity [33] and encompasses two Rossmann-like FAD-binding domains (Pfam: Pyr_redox_2) and a C-terminal mixed alpha-beta domain (Pfam: Reductase_C) [32].

Upon apoptotic stimuli, AIFM1 is cleaved to produce a C-terminal soluble fragment that is translocated to the nucleus to trigger downstream apoptotic events. AIFM1 is also essential for cell viability as its downregulation results in the loss of CI subunits. The pro-survival activity of AIFM1 is tied to its interaction with CHCHD4, a CHCH domain-containing protein located in the IMS [34]. CHCHD4 (the ortholog of yeast protein Mia40) is a crucial component of the redox-regulated MIA machinery responsible for the import of a set of cysteine-containing protein substrates, such as COX17, COX19, MICU1 and COA7 [35]. CHCHD4 functions as a chaperone and catalyzes the formation of disulfide bonds in these substrate proteins, including some respiratory chain subunits and proteins with a variety of other functions such as redox regulation and antioxidant response [35].

We identified CHCHD4-AIFM1 as one of the top scoring PPI pairs with both AlphfaFold2 and RoseTTAFold contact probabilities above 0.99. The AlphaFold2-predicted complex structure between them is shown in Figure 2A. Their interaction is primarily mediated by the N-terminus of CHCHD4, which is missing in the solution structure of CHCHD4 and could be disordered on its own [36]. The predicted interacting residues of CHCHD4 are not from the CHCH domain (residue range: 45 to 109) with two (CX_9_C)_2_ motifs [37, 38]. Instead, our predicted interaction site is consistent with experimental studies describing the N-terminal CHCHD4 27-residue segment being sufficient for its interaction with AIFM1 [34]. Our prediction also mapped the interaction surface on AIFM1, which has eluded experimental studies thus far. The AIFM1 surface is mainly on the edge β-strand (residue 504-508) in the C-terminal Reductase_C domain (Figure 2A). The N-terminal segment of CHCHD4 forms a β-hairpin that interacts with this edge β-strand (Figure 2A). More than 20 missense mutations in AIFM1 (magenta positions in Figure 2A) have been reported in association with several diseases and have been classified as pathogenic/likely pathogenic [39-42]. Interestingly, one likely pathogenic mutation from AIFM1 (Y560H) is mapped near the interaction site (magenta sidechain shown in Figure 2A), and it is predicted to interact with two hydrophobic residues (I12 and F14) on CHCHD4 (sidechains shown in green spheres in Figure 2A). Interactions between these hydrophobic residues could contribute to the binding energy between CHCHD4 and AIFM1 and could explain potential deleterious effects for this mutation.

**Figure 2.**
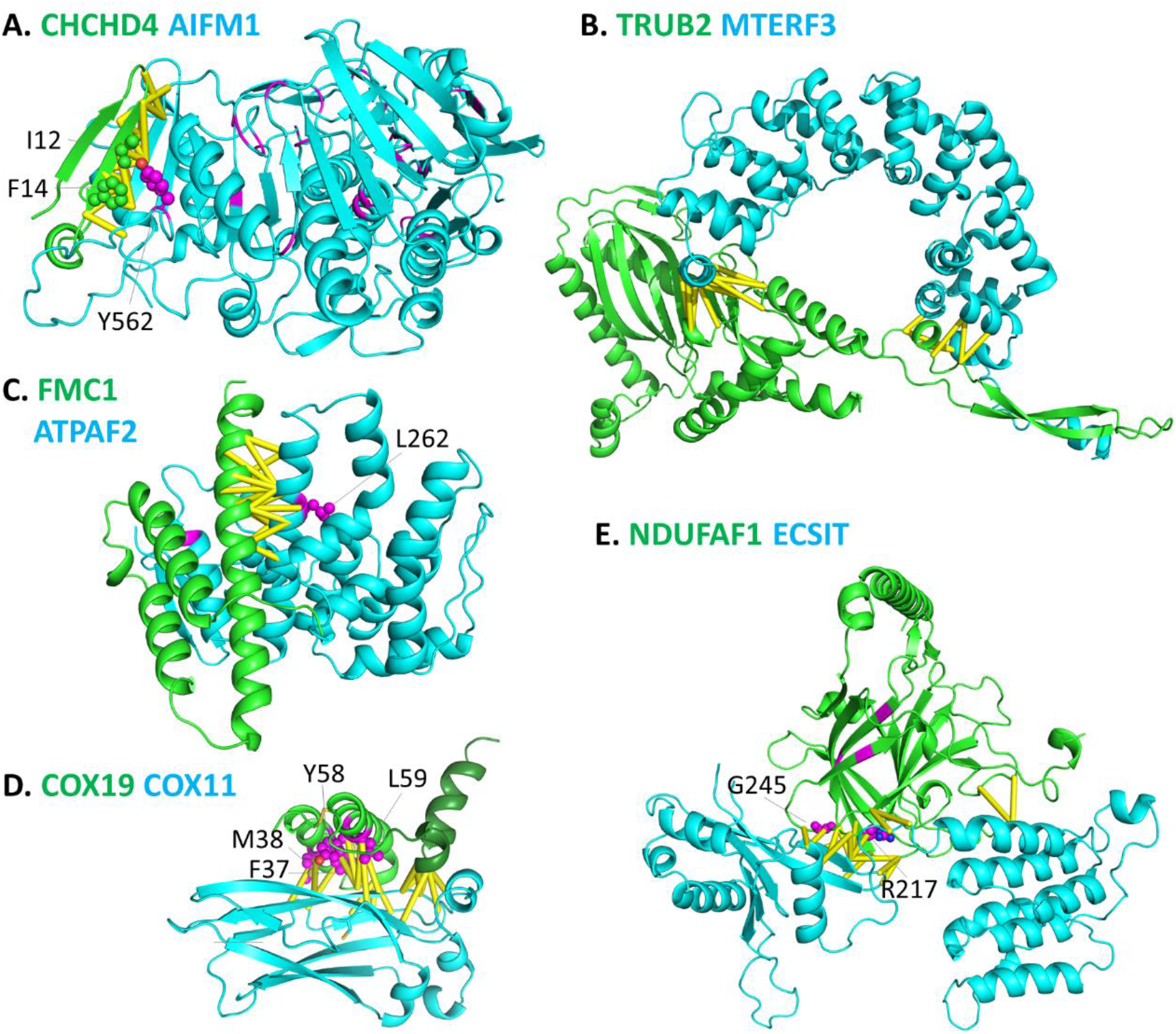
AlphaFold2 structural models of protein complexes without experimentally determined 3D structures. **A**. CHCHD4-AIFM1. Residues involving missense disease-causing mutations are colored in magenta, and those near the interaction interface have their sidechains shown in spheres. Sidechains of the two hydrophobic residues of CHCHD4 interacting with AIFM1 Y562 are shown in green spheres. **B**. TRUB2-MTERF3. **C**. FMC1-ATPAF2. Residues involving missense disease-causing mutations are colored in magenta, and those near the interaction interface have their sidechains shown in spheres **D**. COX19-COX11. Sidechains of the YL and FM signature sequences are shown in magenta spheres. The CHCH domain is shown in light green and the C-terminal two helices are shown in dark green. **E**. NDUFAF1-ECSIT. Residues of missense disease-causing mutations are colored in magenta, and those near the interaction interface are shown in spheres.

#### MTERF3 and TRUB2

MTERF3 (mitochondrial transcription termination factor 3, also named mitochondrial transcription termination domain containing 1 (MTERFD1)) resides in the mitochondrial matrix and is a negative regulator of mitochondrial DNA transcription [43]. MTERF3 also regulates mitochondrial ribosome biogenesis in vertebrates [44] and is a modular factor in mitochondrial protein synthesis [45]. The crystal structure of human MTERF3 revealed that it adopts a right-handed superhelix fold with an overall shape of a half donut [46]. The superhelix consists of tandem repeats of three α-helices named the MTERF-motif [46]. It was proposed that MTERF3 could interact with DNA using the concave side of the half-doughnut that harbors positively charged residues [46].

Recently, MTERF3 was found to colocalize with other proteins in mitochondrial RNA granules, which play crucial roles in post-transcriptional modifications of mitochondrial RNAs and the assembly of the mitochondrial ribosome [47, 48]. MTERF3 is a component of the multi-functional pseudouridine synthase module [48] within the RNA granule together with enzyme components responsible for pseudouridylation of 16S rRNA and mitochondrial mRNAs. TRUB2 is such a pseudouridine synthase that has been shown to interact with MTERF3, and depletion of MTERF3 led to significant decrease in TRUB2 as well as other components of the module [48]. Our coevolution analysis yields high contact probability scores (>0.99) by both AlphaFold2 and RoseTTAFold for MTERF3 and TRUB2, consistent with the experimentally verified physical interaction between them. Furthermore, the structure model of MTERF3 with TRUB2 sheds light on the interaction mode. Interestingly, coevolving residue pairs form two potential interaction sites involving distal ends of the MTERF3 superhelix domain (Figure 2B). Such interactions convert the half-donut shape of MTERF3 to a ring-like complex structure with TRUB2 that maintains an empty space in the middle. This empty space could accommodate the binding of RNA molecules in the RNA granule. Such a structure could be important for maintaining the pseudouridine synthase module within the RNA granule and could play a role in delivering RNA substrates to the pseudouridine synthase, TRUB2, for efficient reactions of pseudouridylation.

#### FMC1 and ATPAF2

A functional connection between human mitochondrial proteins FMC1 (formerly named C7orf55) and ATPAF2 (ATP synthase F1 complex assembly factor 2) was recently predicted by a computational tool (CLustering by Inferred Co-expression, CLIC) [49]. The interaction between these two proteins was also experimentally validated [49]. FMC1 belongs to the LYRM family of proteins [50, 51], and ATPAF2 is one of the assembly factors of CV [52]. This interaction was also found in two high-throughput PPI studies (reported in BioGRID) [14, 16]. Our PPI screen ranked the FMC1 and ATPAF2 pair among the top hits with both AlphaFold2 and RoseTTAFold contact probabilities above 0.99. The FMC1 protein is predicted to adopt a three-helix bundle fold, typical of the LYRM family proteins. The interaction site on FMC1 is mapped to the third core α-helix of such a fold (Figure 2C). The structure model of ATPAF2 contains a small 3-stranded β-sheet at the N-terminus, while the rest of the protein is mainly α-helical. The interaction site on ATPAF2 is mapped to the last α-helix and includes one residue (magenta sidechain shown in spheres in Figure 2C) with a likely pathogenic mutation (L262P) reported in the ClinVar database [41].

#### COX19 and COX11

COX19 is a conserved protein found in mitochondria of a diverse range of eukaryotic organisms including fungi, plants, and animals [53]. Like CHCHD4, COX19 belongs to a family of IMS proteins that contain the CHCH domain with a pair of CX9C signatures [37, 38]. The conserved cysteines in the CX9C motifs form two disulfide bonds in a pair of short antiparallel α-helices. Recently, COX19 was found to interact with COX11, a single-pass transmembrane protein involved in copper transfer to the Cu_B_ center of the COX1 subunit in the cytochrome c oxidase complex [54]. The interaction with Cox11 is required for maintaining stable levels of Cox19 in mitochondria [54]. Both COX19 and COX11 are classified as copper chaperones and CIV assembly factors [17, 55].

Our analysis revealed strong AlphaFold2 and RoseTTAFold contact probability scores (>0.99) between COX19 and COX11. Predicted interacting residues on COX19 are mainly mapped to the α-helical hairpin of the CHCH domain with the twin CX9C motifs (Figure 2D). The CX9C motifs in COX19 proteins are characterized by two conserved YL diads, giving rise to the refined motif signatures of Cx_6_YLxC [54]. In the yeast system, these YL diads were proved to be crucial for the binding to COX11 based on mutagenesis studies [54]. This binding mode is consistent with our model, which showed high contact probabilities for residues corresponding to yeast YL diads in the human COX19 protein, where the first YL diad is changed to FM and the second YL is maintained (magenta residues in Figure 2D). The Cx_6_YLxC α-hairpin of COX19 mainly interacts with the second, third, and sixth core β-strands of the immunoglobulin domain of COX11 [56] (Figure 2D). The COX19 structural model also includes two small α-helices (colored in dark green in Figure 2D) C-terminal to the Cx_6_YLxC helix hairpin (colored in lighter green). Residues in the turn between these two C-terminal α-helices are also predicted to interact with COX11 residues from the loop before the first core β-strand and the loop between the second and third β-strands (Figure 2D).

#### ECSIT and NDUFAF1

ECSIT (Evolutionarily Conserved Signaling Intermediate in Toll pathways) was originally identified as a protein involved in the Toll and bone morphogenetic protein signaling pathways [57, 58]. Later studies showed that ECSIT is a CI assembly factor that interacts with other assembly factors such as NDUFAF1 and ACAD9 [59, 60]. The interaction between ECSIT and NDUFAF1 is strongly supported by AlphaFold2 and RoseTTAFold (contact probability >0.99). The AlphaFold2-predicted ECSIT structure has an N-terminal domain consisting of α-helical repeats and a C-terminal alpha+beta domain with α-helices sandwiching a mainly anti-parallel β-sheet of six β-strands (cyan structure in Figure 2E). The AlphaFold2 model of NDUFAF1 has an N-terminal α-helix with mainly hydrophilic residues and a C-terminal β-sandwich with a jelly roll topology (green structure in Figure 2E). Multiple NDUFAF1-interacting sites of were predicted in ECSIT, including the β-hairpin formed by the third and fourth core β-strands of the second domain, the C-terminal end of the first α-helix of the second domain, the last α-helix of the first domain, and the loop in between the two domains. The interaction surface on NDUFAF1 is mainly mapped to the loop regions at one end between the two β-sheets of the jelly roll domain. Two pathogenic mutations responsible for Mitochondrial complex I deficiency, nuclear type 11 (R217C and G245R, colored in magenta in Figure 2E) in NDUFAF1 [61] lie in the predicted interface between NDUFAF1 and ECSIT, and they are predicted to interact with residues in ECSIT with high contact probability.

#### COQ3 and COQ6

COQ3 [62] and COQ6 [63] are two enzymes in the ubiquinone (coenzyme Q, CoQ) synthesis pathway, an essential component of the mitochondrial oxidative phosphorylation machinery. CoQ is synthesized by the combined actions of at least nine proteins (COQ1–9) [64]. COQ6 is a monooxygenase that converts the C5-hydroxylation of 3-decaprenyl-4-hydroxybenzoic acid (DHB) to 3,4-dihydroxy-5-decaprenylbenzoic acid (DHDB) [65]. DHDB is then converted to 3-methoxy-4-hydroxy-5-decaprenylbenzoic acid by the S-adenosyl-L-methionine-dependent methyltransferase COQ3 [62], which adopts a classical Rossmann-like fold [66].

While the interaction between COQ3 and COQ6 was supported by several experimental studies [14, 67, 68], their complex remains undetermined. The COQ3-COQ6 interface modeled by AlphaFold2 is mostly restricted to the two α-helices before and after the first core β-strand in COQ3 and the loop region between two C-terminal long α-helices of COQ6 (Figure 3A). Since COQ6 and COQ3 are involved in consecutive steps in CoQ biosynthesis, their interaction in a protein complex could facilitate the transfer of the product of COQ6 to the active site of COQ3 for efficient catalysis. Several missense mutations in COQ6 have been associated with the autosomal recessive disorder of Primary coenzyme Q10 deficiency-6 (OMIM: 614650) [69]. One of them, R360L (sidechain shown in magenta spheres in Figure 3A), involves an arginine residue near the interaction interface. In the structure model, R360 forms a salt bridge with D402 of COQ6 that showed high contact probability with a couple of residues from COQ3 including the positively charged residue R166. The presence of these charged residues among others (such as K202 from COQ6 and D191 from COQ3) in the predicted interface highlight the importance of electrostatic interactions in the COQ3-COG6 complex and elucidates a possible molecular mechanism for Primary coenzyme Q10 deficiency-6.

**Figure 3.**
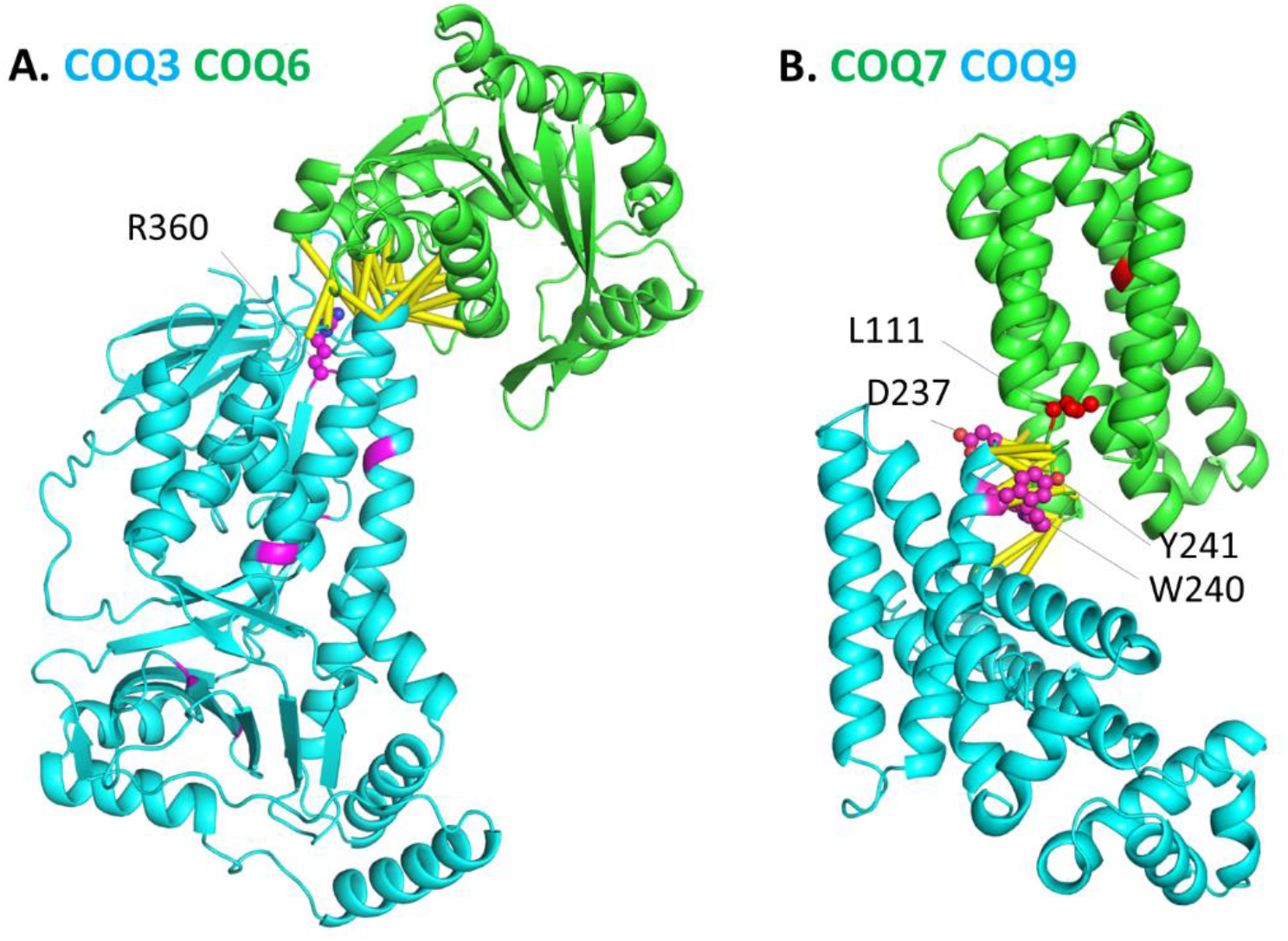
AlphaFold2 structural models of protein complexes in the ubiquinone biosynthesis pathway. **A**. COQ3-COQ6. Missense disease-causing mutations in COQ3 are colored in magenta, and the sidechain of R360 near the interaction interface is shown in spheres. **B**. COQ7-COQ9. Residues involving missense disease-causing mutations in COQ7 are colored in red, and sidechain of L111 near the interaction interface is shown in spheres. Sidechains are shown in magenta spheres for COQ9 residues whose mutagenesis affected the binding to COQ7.

#### COQ7 and COQ9

COQ7 is an oxidoreductase that catalyzes the hydroxylation of 2-polyprenyl-3-methyl-6-methoxy-1,4-benzoquinol to form 3-demethylubiquinol, the penultimate step in CoQ biosynthesis [70]. While the structures of human COQ7 and its close homologs have not been solved, HHpred similarity searches suggest that it is evolutionarily related to a large family of heme oxygenases and ribonucleotide reductases with a Ferritin-like helix-bundle fold [70]. AlphaFold2 predicts COQ7 to adopt a helix-bundle fold consisting of six α-helices, consistent with the HHpred results. The lipid-binding protein COQ9 functions in CoQ biosynthesis through its interaction with COQ7 and the stabilization of the whole CoQ biosynthetic complex [71].

Recent structural studies on COQ9 revealed that it adopts an ancient fold of the bacterial TetR family transcriptional regulators [71]. While the structure of the COQ7-COQ9 complex has not been determined, mutagenesis studies suggested that the interaction site on COQ9 corresponds to a conserved surface patch comprising mostly of the region between the seventh and eighth α-helices [71]. Such experimental results are consistent with our predictions, which mapped the interacting residues from COQ9 to this surface patch (Figure 3B). Several COQ9 residues, such as D237, Y240, and W241 (sidechains shown as magenta spheres in Figure 3B), show high contact probabilities with residues of COQ7. Their mutations (W240K, W240D, Y241K, and D237K) maintained normal melting temperature of

COQ9, while they abolished the interaction with COQ7 [71]. Our coevolution analysis further suggests the interaction site of COQ7 to be the C-terminal end of the second core α-helix and the loop region between the second and third core α-helices (Figure 3B). Missense mutations in COQ7 have been associated with the disorder of Coenzyme Q10 deficiency, primary 8 (OMIM: 616733) [72, 73]. One such mutation L111P (sidechain shown as red spheres in Figure 3B) is mapped near the interaction interface of COQ7 and COQ9, suggesting that its deleterious effect could be related to the destabilization of the COQ7-COQ9 complex.

#### Other protein complexes supported by coevolution analysis

Besides the above examples, we identified several high-scoring PPIs supported by experiments. They include the mitochondrial pyruvate carrier complex, a heterodimeric transporter formed by MPC1 and MPC2 [74, 75]. MPC1 and MPC2 are distantly related proteins with three transmembrane segments. HHpred results suggest that MPC proteins (Pfam: MPC) are remotely related to PQ-loop family of sugar transporters [76], and they have been classified in the same Pfam clan (MtN3-like). Consistent with this classification, the AlphaFold2 models of MPC1 and MPC2 adopt the same fold as the PQ-loop transporters [76, 77]. In addition, the right-handed three transmembrane segments of MPC1 and MPC2 form a pseudosymmetric dimer arranged in the same fashion as PQ-loop transporters (Figure 4A). MPC1 and MPC2 exhibit high contact probability mainly between the second transmembrane segment of MPC1 and the first transmembrane segment of MPC2 (Figure 4A). Two other predicted PPIs with contacting residues mainly from transmembrane segments are the pair of COX16 and SCO1 (both CIV assembly factors) [78] (Figure 4B) and the pair of HIGD2A (a CIV assembly factor) and MT-CO3 (a core subunit of CIV) [79, 80] (Figure 4C).

**Figure 4.**
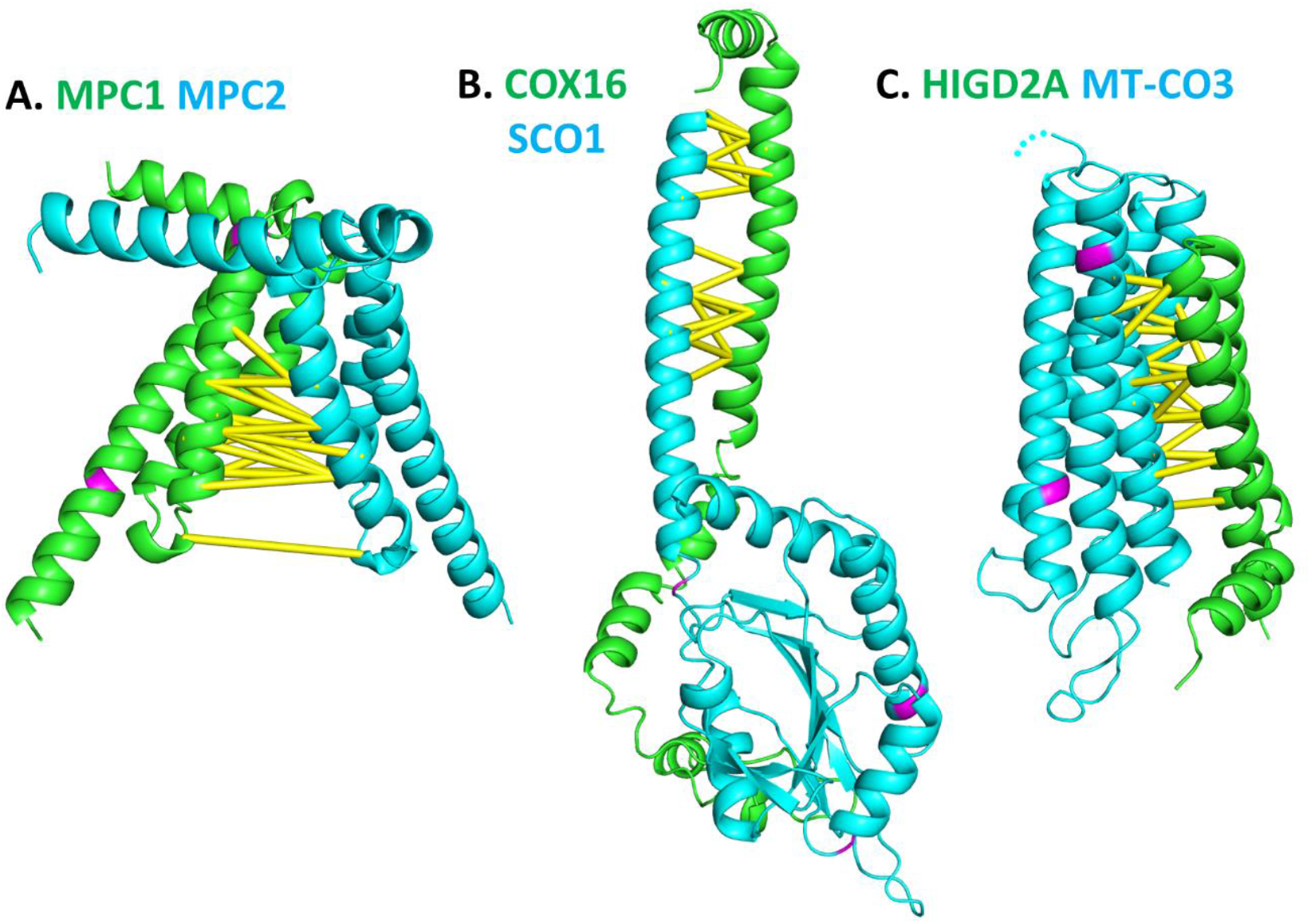
AlphaFold2 structural models of protein complexes with predicted contacts in transmembrane segments. **A**. MPC1-MPC2. **B**. COX16-SCO1. **C**. HIGD2A-MT-CO3. Residues involving missense disease-causing mutations are colored in magenta, and those near the interaction interface have their sidechains shown in spheres.

### Coevolution analysis identified potential interaction partners for poorly characterized mitochondrial proteins

About 20% of mitochondrial proteins are uncharacterized and do not have clear function annotations [14]. Studies on PPIs by experimental or computational methods can help elucidate their cellular functions [49], as elucidated by the following examples.

#### Identification of PYURF as a potential subunit of the NDUFAF5 hydroxylase complex

Affinity enrichment mass spectrometry (AE-MS) was used in a previous study to identify PPI of 50 select mitochondrial uncharacterized proteins (MXPs) [14]. One MXP, formerly named C17orf89, was found to interact with NDUFAF5 and play an important role in CI assembly [14]. This protein has since been renamed NDUFAF8. We identified the NDUFAF8-NDUFAF5 complex in our screen with the contact probability of 0.911 by RoseTTAFold and the contact probability of 0.444 by AlphaFold2. We also identified another MXP named PYURF that likely interact with NDUFAF5. The PYURF-NDUFAF5 pair has a AlphaFold2 score of 0.998 and a RoseTTAFold score of 0.959. Predicted interaction with NDUFAF5 suggests that the function of the small uncharacterized protein PYURF could be related to CI assembly as well.

NDUFAF5 adopts a Rossmann-like fold with seven β-strands and belongs to the family of S-adenosylmethionine-dependent methyltransferases [81]. However, instead of being a methyltransferase, it catalyzes the hydroxylation of an arginine residue in of the CI subunit NDUFS7, and this posttranslational modification is crucial in the early stage of CI assembly [81]. As an essential assembly factor, NDUFAF5 has mutations causing the disease of mitochondrial complex I deficiency (nuclear type 16, OMIM: 618238) [82]. Mapping of the predicted interface residues onto the structure models of NDUFAF8, NDUFAF5 and PYURF suggests that NDUFAF8 and PYURF occupy distinct interaction sites on NDUFAF5 (Figure 5A) that are distal from each other. NDUFAF5 (cyan in Figure 5A) residues coevolving with NDUFAF8 (orange in Figure 5A) is mapped to an α-helix before the last β-strand and is far from the active site. A likely pathogenic mutation (F55L, sidechain of F55 shown in blue spheres in Figure 5A) in NDUFAF8 [83] was mapped near its interaction site with NDUFAF5.

**Figure 5.**
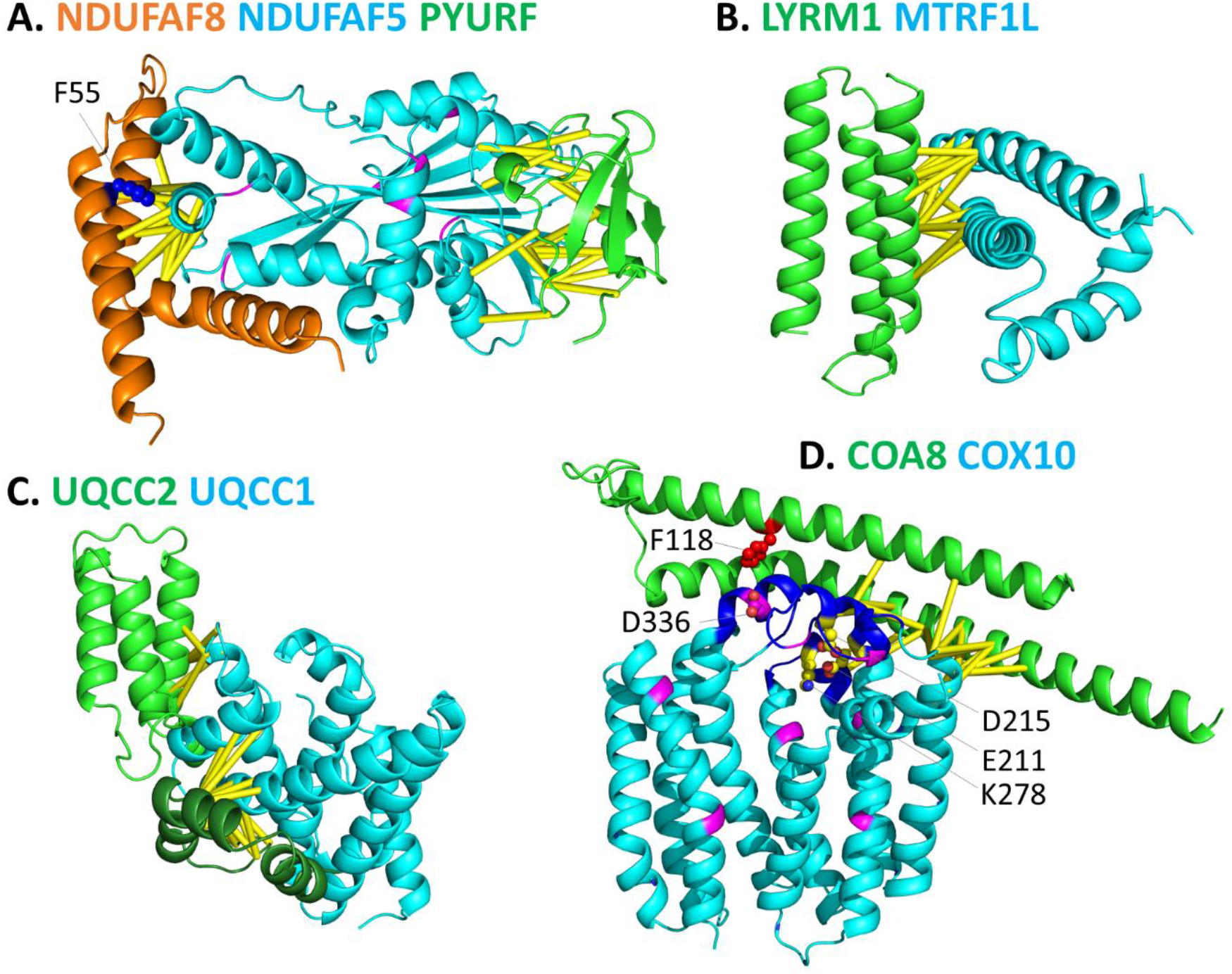
AlphaFold2 structural models of protein complexes involving proteins with poorly characterized proteins. **A**. NDUFAF8-NDUFAF5-PYURF. Residues involving missense disease-causing mutations of NDUFAF5 are colored in magenta. Sidechain of NDUFAF8 F55 involving a missense disease-causing mutation is shown in blue spheres. **B**. LYRM1-MTRF1L. **C**. UQCC1-UQCC2. **D**. COA8-COX10. The CAP domains of COX10 are colored in blue. COX10 residues involving missense disease-causing mutations are colored in magenta, and those near the interaction interface have their sidechains shown in spheres. D215, E211, and K278 of COX10 (with carbon atoms colored in yellow) are conserved charged residues likely involved in catalysis. The residue (F118) involving a missense disease-causing mutation in COA8 has its sidechain shown in red spheres.

NDUFAF5 residues coevolving with PYURF (green in Figure 5A) are mapped near the active site to an edge β-strand, the α-helix before it and the Rossmann crossover loop after it. PYURF is a small protein predicted to adopt a Trm112p-like fold [84]. Remote homologs of PYURF include Rieske-like ferredoxins and the Trm112 proteins in archaea and eukaryotes [85]. Interestingly, Trm112 proteins are subunits of multifunctional methyltransferases that form complexes with and activate the tRNA methyltransferases such as Trm11 and Trm9. Structures of the recently solved archaeal Trm112-Trm11 complexes [86, 87] revealed similar interaction sites as predicted by AlphaFold2 in PYURF and NDUFAF5. Our coevolution analysis as well as the remote homology of PYURF to Trm112 proteins suggests that PYURF is a subunit that forms a complex with the NDUFAF5 hydroxylase and could play an essential role in the catalytic activity. The involvement of PYURF in CI assembly was supported by a recent genome-wide CRISPR death screen that shows the phenotype of CI deficiency when the PYURF gene was disrupted [88].

#### Identification of potential interaction partners and interfaces of LYRM proteins

The LYRM (leucine/tyrosine/arginine motif) family proteins (also called LYR proteins) are small proteins with diverse functions. LYRM proteins are found to associate with OXPHOS complexes as CI subunits (NDUFB9 (LYRM3) and NDUFA6 (LYRM6)), CI assembly factor (LYRM2) [89], CII assembly factors (SDHAF1 (LYRM8) and SDHAF3 (LYRM10)) [90], CIII assembly factor (LYRM7) [91], and CV assembly factor (FMC1) [49]. Other LYRM proteins are involved in Fe–S cluster biosynthesis (LYRM4, also known as ISD11) [89], mitoribosome assembly and translation (AltMIEF1 (also named AltMiD51 or L0R8F8)) [92-94], and regulation of the electron transferring flavoprotein complex (LYRM5) [50, 51]. LYRM proteins contain the LYR domain that adopts a three-helix bundle fold [93]. Our top predictions include several known interactions involving LYRM proteins, such as FMC1-ATPAF2 (described above), LYRM4-NSF1 that is important for Fe–S cluster biosynthesis, NDUFB9/LYRM3-NDUFB3 in CI, and NDUFA5-NDUFS3 in CI. We identified NDUFA5 (Pfam: ETC_C1_NDUFA5), an accessory subunit of CI, as a distant member of the LYRM family. Remote homology between NDUFAF5 and other LYRM proteins is supported by HHpred search results, the presence of LYR motif in NDUFA5, and the three-helix bundle fold of NDUFA5 in CI structures [95, 96].

The functions of LYRM1 and LYRM9, two LYR-domain containing proteins, remain to be elucidated [94, 97]. In our PPI screen LYRM1 is predicted to interact with MTRF1L (mitochondrial translational release factor 1-like) (AlphaFold2 score: 0.902, RoseTTAFold score: 0.955). MTRF1L is responsible for translation termination upon detection of certain termination codons [93]. In the structural model of the LYRM1-MTRF1L complex, the interface residues in LYRM1 are mapped to the first α-helix of the LYR domain with a three-helix bundle fold. The interface residues from MTRF1L are mapped to a N-terminal α-helical domain [98] (Figure 5B). RoseTTAFold also reported modestly high contact probability scores for the LYRM1-LYRM9 pair (score: 0.862) and the LYRM9-MRPL57 (mitochondrial ribosomal protein 63) pair (score: 0.907), albeit the AlphaFold2 scores for them are low (below 0.2). AltMIEF1, a small protein encoded by an open reading frame in the 5′ region of the MIEF1 gene, was the only LYRM protein previously found to function in translation and ribosomal assembly [92]. Our study suggests that the uncharacterized LYRM1 and LYRM9 could also play important roles in regulation of translation and other functions related to mitochondrial ribosomes.

HHpred sequence similarity searches and Alphafold2 structural models also suggest the presence of a divergent LYR domain in UQCC2, a small protein that interacts with UQCC1 and functions in the assembly of CIII [99]. UQCC1 and UQCC2 are orthologs to fungal CIII assembly factors Cbp3p and Cbp6p, respectively. AlphaFold2 structural model predicts that UQCC1 adopts an α-helical bundle fold with six major α-helices, consistent with the experimental structure of its bacterial homolog [100]. Our analysis predicted high probability for interaction between UQCC1 and UQCC2. The predicted UQCC1-UQCC2 interface in UQCC2 is mapped to the N-terminal LYR domain (colored in lighter green in Figure 5C) that adopts a three-helix bundle fold, as well as regions C-terminal (colored in darker green in Figure 5C) to the LYR domain. Our findings suggest UQCC2 as a new member of the LYRM family that functions in CIII assembly [101].

#### Identification of COA8 as a potential subunit of the COX10 enzyme complex

COA8, previously named APOPT1 (apoptogenic-1), was recently discovered to function in CIV assembly. Loss-of-function mutations in COA8 [102, 103] caused the disease phenotype of leukoencephalopathy associated with mitochondrial cytochrome *c* oxidase (COX) deficiency, and gene knockout of COA8 in mouse [104] and knockdown in *Drosophila* [105] resulted in reduced cytochrome c oxidase activity and levels. The functional mechanism of COA8 in CIV assembly has not been revealed. Our analysis identified high contact probability scores (>0.99 by both AlphaFold2 and RoseTTAFold) between COA8 and COX10. As an essential CIV assembly factor, COX10 catalyzes the farnesylation of the vinyl group of heme B to produce heme O, the first step of the mitochondrial heme A biosynthesis required for CIV assembly.

The predicted interaction between COA8 and COX10 suggests that COA8 could be a subunit of the heme farnesyltransferase complex. The AlphFold2 model of the COA8-COX10 complex revealed that COA8 forms two long α-helices, and COX10 is a multi-pass transmembrane protein with 9 transmembrane segments, consistent with its homology to proteins in the UbiA prenyltransferase family (Pfam: UbiA) [106, 107] (Figure 5D). The predicted interface residues are mainly mapped to the middle of the second α-helix in COA8 and in between the second and third transmembrane segments of COX10 (Figure 5D), which is part of the cap domains (colored in blue in Figure 5D) previously defined as the connections between transmembrane helices near the active site [107]. Several missense mutations of COX10 have been reported in mitochondrial disorders characterized by cytochrome c oxidase deficiency. Most of the pathogenic mutations in COX10 (colored magenta) are not close to the predicted interface between COA8 and COX10. Most of the pathogenic mutations in COA8 resulted in loss-of-function by introducing premature stop codons or frame shifts [108]. One recently found missense pathogenic mutation in COA8, F118S [103] (sidechain shown in red spheres in Figure 5D), lies in the predicted interface of the COA8-COX10 complex and is close to residue D336 in COX10 that also harbors pathogenetic mutations (D336G and D336V) [109]. Our results suggest that those pathogenetic mutations may cause disease phenotypes by disrupting the proper interaction between COA8 and COX10.

## Materials and methods

### Generation of protein sequence alignments

The eukaryotic proteomes were downloaded from the NCBI genome database. It consists of 49102568 proteins from 2568 representative or reference genomes. The list of human mitochondrial proteins was obtained from the MitoCarta3.0 database [17]and their sequences were obtained from the UniProt database [42]. For each human mitochondrial protein (called a target protein), the corresponding orthologous group at the eukaryotes level in OrthoDB [110] was identified. This group of orthologous proteins were clustered by CD-HIT [111] at 40% identity level. For each CD-HIT cluster, we selected one representative sequence that showed the best BLAST [112] score to the target protein. The representative sequences were then used as queries to search against the NCBI eukaryotic proteomes to identify homologous proteins to the target protein.

For each organism we identified up to one protein (best hit, if available) that shows the highest sequence similarity to the target protein from the combined BLAST hits found by multiple representative sequences. These best hits together with the target protein are considered as the expanded orthologous group for the target protein. This group of sequences were subject to multiple sequence alignment by MAFFT (with the --auto option) [113]. To construct the joint alignment for any two target proteins, we identified the intersection of the organisms for their orthologous groups and merged the sub-alignments containing only these organisms for the two proteins. Positions that are gaps in the human proteins were removed. Predictions of coevolving residues by deep learning neural network were carried out for protein pairs that have a combined length of less than 1500 amino acid residues.

### Selection of top predictions

We applied the RoseTTAFold 2-track model, a faster but inferior model than the 3-track one, to each concatenated alignment [ref]. RoseTTAFold predicts the probability density for the distance between each residue pair. We summed up the probability for distance bins below 10Å and used it as the contact probability (ranging from 0 to 1) of a residue pair. Residue pairs involving the last 10 residues in the first sequence and the first 10 residues of the second sequence were discarded as they represent artificially continuous segment in the merged alignment and could have biased scores. To further improve the performance, we also removed residue pairs involving the mitochondrial target sequences (defined in the UniProt Feature fields) in either protein. The best contact probability of the remaining residue pairs was reported as the contact probability score for the protein pair.

We observed that some proteins enriched in top hits with high contact probability scores could be false positives. To downgrade the influence of these proteins, we designed a score to penalize proteins involving many high-scoring pairs. For each protein, we recorded up to 20 contact probability scores of its highest scoring pairs with contact probability scores more than 0.5. If the number of pairs (*N*) with scores more than 0.5 for a protein is less than 20, we extended the list of its top scores by adding 20 − *N* pseudo scores of 0.5 at the end, such that the list consists of 20 numbers. The average of the 20 numbers in this score list is then used a penalty weight, and its minimum value is 0.5. The new adjusted score for any protein pair is calculated as the original score divided by the maximum of the two penalty weights of the two proteins.

We selected the top 200 pairs ranked by the original contact probability score and the top 350 pairs ranked by the adjusted score. Their union consists of 401 pairs of proteins. AlphaFold2 was used to build structural models for these 401 pairs. AlphaFold2 also produces probability distribution for residue-residue distances, and we computed contact probability as the sum of probability for distance bins below 10 Å. The highest contact probability between residues of two proteins was used to indicate the contact probability for this protein pair.

### Performance evaluation

For precision-recall analysis, we excluded protein pairs that are potentially paralogs (defined as sequence identity >40%) as they could have artificially high contact scores by sharing sequences in their alignments. For the rest of the pairs, we considered a pair of proteins to be positive cases of PPI if it has been reported by three or more experimental studies in the BioGRID database, or if both proteins have significant sequence similarity (>30% sequence identity) to two interacting chains in an experimentally determined structure of a protein complex. A total of 1563 positive pairs (473 supported by BioGRID and 1193 supported by experimental complexes) were found in our dataset. Negative cases of PPIs are those protein pairs without experimental structures and were never reported in the BioGRID database. For precision-recall analysis, we used the set of positive pairs and a randomly selected subset of negative pairs that are 100 times more than the positive pairs.

For the PPIs supported by known experimental complex structures, we evaluated the agreement of top-scoring residue pairs provided by RoseTTAFold against the experimentally determined interface residues. BLAST was used to detect proteins with known structures for each human mitochondrial protein. The BLAST alignment was used to map the residues in a target protein to those residues in its homologs with known structures. A residue pair predicted by RoseTTAFold (defined as those with contact probability >0.5) is considered to lie in the interaction interface of an experimental complex if the corresponding aligned residue pair in the complex structure has an inter-residue non-hydrogen atom pair within 10Å. The fraction of true interacting residue pairs was then calculated as the number of residue pairs lying in the experimental interface divided by the total number of residue pairs reported by RoseTTAFold.

## Supplementary materials

**Supplementary Table S1**. RoseTTAFold and AlphaFold2 scores of top-scoring protein pairs by RoseTTAFold.

**Supplementary Table S2**. RoseTTAFold and AlphaFold2 scores for protein pairs with BioGRID interaction count ≥ 3.

## Acknowledgement

We would like to thank Minkyung Beak for sharing the RoseTTAFold script prior to its publication, Aaron Guillory for help in maintaining the computer cluster used in this study, Lisa N. Kinch and Nick V. Grishin for helpful discussions and suggestions. Qian Cong is a Southwestern Medical Foundation scholar.

